# A Rapid Method for Producing Adeno-Associated Viral Vectors Suitable for Transducing Rodent Neurons *in vitro* and *in vivo*

**DOI:** 10.1101/2024.05.06.591977

**Authors:** Doug B. Howard, Reinis Svarcbahs, Lana N. Gore, Brandon K. Harvey, Christopher T. Richie

**Affiliations:** National Institute on Drug Abuse Intramural Research Program Genetic Engineering and Viral Vector Core; National Institute on Drug Abuse Intramural Research Program

**Keywords:** Adeno-associated viral vectors, AAV, neuroscience, viral packaging

## Abstract

The use of adeno-associated viral vectors for delivery of genetic information into the mammalian CNS remains popular but producing highly purified vectors for in vivo applications requires a significant investment of resources and time that can impede the development and testing of AAV vectors for experimentation. To address this issue, we have developed a simplified AAV packaging protocol that does not require large capital equipment (ultracentrifugation or chromatography machines) yet still produces virus in quantities that are sufficient for testing AAV prototypes in the rodent CNS. This protocol is serotype agnostic, and has been successful with AAV1, AAV9, AAV-DJ, and rAAV2-retro. Intracranial injection of AAV-EF1a-GFP-KASH into rats demonstrated that our “small scale” AAV preps produce patterns of transgene expression and inflammation that are similar to those produced by the same AAV vector purified by affinity column chromatography. Our protocol allows for multiple vectors to be packaged and processed in parallel, making it ideal for testing multiple variants, constructs, and prototypes simultaneously.

## INTRODUCTION

Adeno-associated viral (AAV) vectors have been used for gene delivery and gene therapy applications for several decades and the demand for this technology continues to grow. Despite being a well-established technology, the high production cost (time and money) relative to other gene delivery modalities (lentiviral, lipid nanoparticles, etc.) limits the development and higher throughput testing of multi AAV vectors for experimentation. Numerous methods for AAV production have been published since the first reports of “helper-free” production of AAV particles by Xiao et al and the purification of AAV by column chromatography by Zolotukhin et al (Xiao et al., 1998; Zolotukhin et al., 1999). Many of these methods make use of a triple-transfection method, whereby HEK293 cells are transfected with three plasmids, encoding the viral vector genome, the AAV replication and capsid genes, and the adenovirus genes necessary for AAV production. The resulting AAV particles are purified from the transfected material by a series of filtration, centrifugation, and/or chromatography steps (Chen et al., 2019; Lock et al., 2010; Sandoval et al., 2019). Although high-quality preps are achievable with ultracentrifugation on a density gradient or affinity column chromatography, the reagents, equipment, and technical workflow are not conducive to high throughput or cost-effective production of small batch sizes suitable for prototyping and pilot studies.

Herein, we describe a method to produce small batches of AAV for applications in rodent CNS. The AAV produced by our method are of sufficient titer and quality to express transgenes in in vitro and in vivo, while bypassing the need for ultracentrifugation and affinity purification.

## METHODS

### Large scale AAV Packaging

Large-scale AAV packaging is performed using a protocol published in Howard, 2008 with some modifications (Sup Doc 1) (Howard et al., 2008). HEK293 cells were seeded onto a 150mm dish and triple-transfected with a mix containing helper plasmid, rep-cap plasmid, and viral vector genome plasmid, using calcium phosphate precipitation. Forty hours post-transfection, the cells were lysed by three freeze/thaw cycles and then treated with SAN HQ nuclease. The nuclease-treated lysate was then passed through a series of filters (5um, 0.45um, and 0.22um) before being loaded onto an AKTA FPLC machine configured with an AVB column or POROS AAV9 column. Virus was eluted, dialyzed, and aliquoted, then stored at -80C until use.

### Small scale AAV Packaging

Small-scale AAV packaging is performed using a scaled down version of the protocol published in Howard, 2008 with some modifications (Sup Doc 2) (Howard et al., 2008). HEK293 cells were seeded onto a single 150mm dish and triple-transfected with a mix containing helper plasmid, rep-cap plasmid, and viral vector genome plasmid, using calcium phosphate precipitation. Forty hours post-transfection, the cells were lysed by three freeze/thaw cycles and then treated with SAN HQ nuclease. The nuclease-treated lysate was then passed through a series of syringe filters (5um, 0.45um, and 0.22um) before being loaded onto 100,000 MWCO Amicon Ultra-15 centrifugal filters and subjected to volume reduction. The sample then underwent three rounds of buffer exchange with dialysis buffer. The final vector-containing retentate (∼150 uL) was then aliquoted and stored at -80C until use.

### Protein electrophoresis and staining

AAV samples were transferred to an Eppendorf Lo-Bind tube and were mixed with ¼ volume of 4xLDS loading dye (Life technologies) containing 2-mercaptoethanol (16 uL total). Samples were heated to 70C for 10 minutes with 650rpm shaking in an Eppendorf thermomixer before loading onto a 4-20%, 10-well Bis-Tris gel (Novex, Life Technologies) with 1X MOPS buffer (Life Technologies) and run for 50 minutes at 200V. Once the gel run was complete, the gel was removed from its plastic casing and stained with SimplyBlue (Invitrogen) using the vendor’s “microwave” method. The stained gel was imaged on an Azure Sapphire Biomolecular Imager.

### Bacterial Contamination testing

Ten microliters of aliquoted virus were added to a single well in a 96-well plate containing 200uL of antibiotic-free tissue culture media (PCN media) and incubated in a humidified incubator at 37C. Wells were monitored daily for sings of bacteria or fungal growth. Lots were deemed “negative for contamination” if there was no growth observed within seven days.

### AAV Titering

The concentration/titer (viral vector genomes (vg) per mL) of each packaged vector was determined by digital PCR with fluorescent probe-based assays. Viral aliquots were diluted (10^-5^, and 10^-6^, and 10^-7^) in PBS, mixed with QIAGEN dPCR mix for probes along with assays for the EF1a promoter or WPRE sequences, and partitioned on 8.5k QIAcuity nanoplates. Primer and probes sequences are listed in (Sup Table 1 – primers and probes).

### Isolation of primary cortical neurons

PCNs were generated using a modified protocol based on Howard et al, 2008 (Howard et al., 2008). Cortices isolated from Sprague Dawley rat E18 embryos (SDECX, BrainBits) were removed from their Hibernate EB solution (HEB, BrainBits), into a 15ml conical tube and rinsed with room-temp PBS, pH 7.4 (# 10010049, Gibco) (1ml/cortex) and centrifugated at 100xg for 5 minutes. The supernatant was then replaced with re prewarmed 0.25% trypsin (# 25200-056, Gibco) (1ml/cortex). Cortices were incubated in trypsin at room temperature for 20 minutes with mild shaking every 5 minutes. Trypsin was then carefully removed, and cortices were gently washed with pre-warmed Neurobasal media (#21103-049, Gibco) three times using twice the trypsin volume per each wash (2ml/cortex). Cortices were then triturated in 5mL of PCN plating media [Neurobasal, 2% Heat Inactivated Fetal Bovine Serum (# F4135, Sigma), 2% B27 (#17504-044, Gibco), 200mM L-glutamine, and 25 mM L-glutamate] by pipetting up and down 10 times using a 5mL serological pipette (# 86.1253.001, Sarstedt). Trituration step was repeated with an additional 5mL of plating media. The final 10mL was then strained through a 100µm strainer (# 352360, Falcon). Cells were counted using trypan blue (# 15250061, Gibco) and the Primary Cell Trypan Blue option on the CellDrop (DeNovix). Cells were plated at 5.00E4 cells per well on the inner 60-wells of a PEI-coated 96-well plate with PBS added to the outer-most wells to minimize evaporation. PCN-seeded plates were incubated at 37°C with 5.5% CO2, and received a 50% media exchange on DIV4, 6, 8, 11, and 13 using PCN maintenance media (Neurobasal, 2% B27, 200mM L-glutamine).

### AAV Transduction in vitro

Viral transductions were performed with the following viruses: AAV1-EF1a-EGFP-KASH (large scale packaging), AAV1-EF1a-EGFP-KASH (small scale packaging), AAV9-EF1a-EGFP-KASH (large scale packaging), and AAV9-EF1a-EGFP-KASH (small scale packaging). All viruses were titer matched and transduced at 2.41E12 VG/mL, 1.00E12 VG/mL, 0.50E12 VG/mL, and 0.25E12 VG/mL. Viral transductions occurred on DIV6 with the removal 50% of the total media volume, followed by the addition of 5ul of virus (diluted to the indicated VG/mL in AAV buffer). The PCNs were incubated at 37°C for 2 hours before 50% volume of fresh media was added to the cells. PCNs were then maintained until DIV13 as described above.

### ATP Assay

The Promega CellTiter-Glo Luminescent Cell Viability Assay (# G7571, Promega) was used to determine cell viability. The CellTiter-Glo Substrate (G755A) was reconstituted in 10mL of the CellTiter-Glo Buffer (G756A) to generate the CellTiter-Glo Reagent. 100uL of media was removed from PCN wells and 100uL of the CellTiter-Glo Reagent was added. The plate was then covered in tinfoil and placed on a plate shaker for 2 minutes. 100uL of the reagent/media mix was removed and plated on an opaque white 96-well plate (# 3917, Corning). The luminescence was then read using the Biotek Synergy H1 plate reader. Analysis was conducted using GraphPad Prism 8 and (ordinary two-way ANOVA).

### Immunofluorescence

PCNs were fixed on DIV15 using 4% paraformaldehyde (PFA) in PBS [pH 7.4] for 20 minutes and then stored in PBS at 4C. Cells were DAPI stained (# D3571, Invitrogen) in a 0.1% Triton (# T8787, Sigma) in PBS solution for 10 minutes at room temperature. Wells were imaged using the EVOS FL Auto 2 (Invitrogen) under the same light, exposure, gain, and objective settings using both a DAPI filter cube (AMEP4650, Invitrogen) and a GFP filter cube (AMEP4951, Invitrogen).

### Animals

Male Long Evans rats were purchased from ENVIGO. Animal studies were approved by the NIDA Animal Care and Use Committees (ACUC) and comply with NIH guidelines for animal research. Rats were group housed under standard laboratory conditions (12 h light/dark cycle; room temperature, 23 ± 2°C; relative humidity, 50 ± 15%).

ASP: 23-INRB-41

### AAV transduction in vivo

Rats were anesthetized with isoflurane (5% induction, 2.5–3% maintenance), and the recombinant AAV1-EF1α-GFP-KASH and AAV9-EF1α-GFP-KASH vectors were bilaterally injected in striatum in a stereotaxic operation. Small-scale packaged and large-scale packaged AAVs were normalized to 2.4E12 VG/mL, and were injected in the opposing hemispheres (volume, 2 μL; rate, 0.5 μL/min) 0.6 mm anterior and 3.0 mm lateral to bregma, and from -5.5 to -4.5 mm below the dura [stereotaxic coordinates according to Paxinos and Watson (Paxinos and Watson, 2007). Rats were returned to home cage post-surgery. Two weeks after AAV delivery, rats were perfused with saline supplemented with heparin (1000 U/mL; #NDC 25021-400-30, Sagent Pharmaceuticals) followed by 4% paraformaldehyde. Brains were removed and transferred to 4% paraformaldehyde for 4 hours followed by 18% and 30% sucrose gradient until equilibrated, as evidenced by sinking to the bottom of tube. Tissue was flash frozen on dry ice and stored at -80°C until further use.

### Immunohistochemistry

Frozen brains were sectioned with a cryostat (Leica, CM1950) at 50 µm. The sections were stained with rabbit anti-Iba1 (1:1500; #019-19741, RRID: AB_839504, FUJIFILM Wako Shibayagi), mouse anti-GFAP (1:2000; #MAB3402, RRID:AB_94844, Millipore). Nuclei were stained with DAPI (4’,6-Diamidino-2-Phenylindole, Dilactate, 1:3000; #D3571, Invitrogen) then mounted on glass slides with Mowiol. Whole brain images were captured by VS200 slide scanner (Olympus, Japan) at 20X. Zoomed in images were taken with a Nikon Eclipse Ti2 inverted confocal microscope at 40X and processed in Fiji-ImageJ.

## RESULTS

### Section 1. Development of small scale protocol

Our standard protocol for AAV production relies on the triple-transfection of HEK293 cells at a scale of 20 x 150 mm dishes, followed by the purification of the viral particles using affinity column chromatography (AVB columns from Cytiva or POROS AAV9 from ThermoFisher).

We analyzed the yields obtained from this workflow over the past twelve years and determined that on average, we obtained 6.0e10 and 2.9e11 viral genomes per 150mm dish for AAV1 and AAV9, respectively (Fig 1A). This historical data is composed of production runs using different viral vector genomes, cell passage numbers, plasmid prep lots, etc. which likely contributes to its apparent variability. We estimated that the number of viral genomes produced per 150 mm would be sufficient for pilot studies or vector validation if it could be concentrated to a volume around 50-100uL.

**Figure 1.**
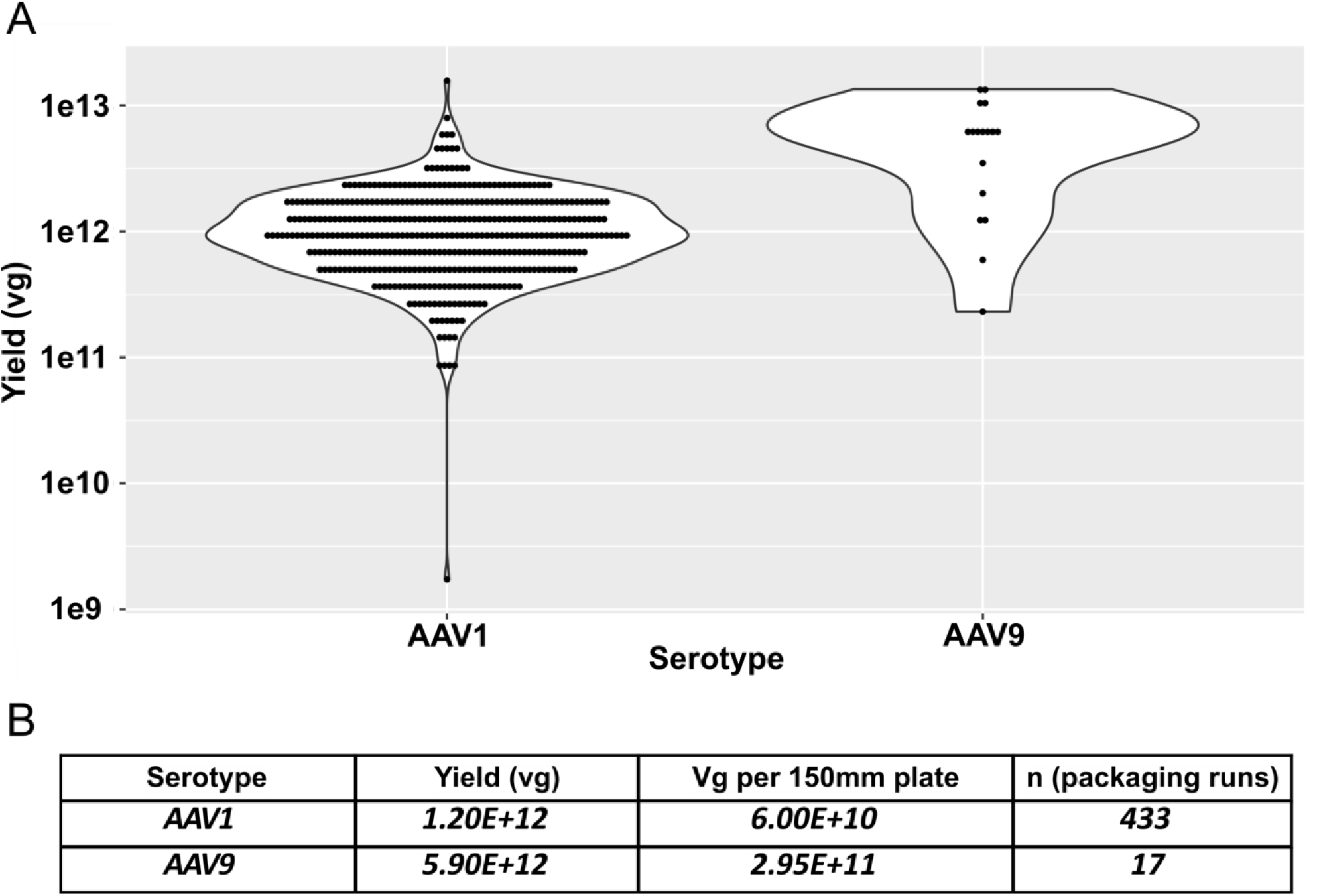
Historical yields from large scale packaging. A) Violin plots showing the yield in total viral particles from our core facility’s large scale (20 x 150 mm plates) packaging for serotypes AAV1 and AAV9 purified by affinity resin. B) the average yield per prep and estimated yield of purified vector obtained per 150mm plate.

We then compiled a cost analysis for the different phases of our AAV production workflow and identified the purification step as the most influential principal component for financial cost, followed by the feedstock plasmids and other consumables (Fig 2). Within the AAV purification step, the most expensive consumable is the pre-packed affinity columns, which has a cost of greater than $300 USD when purchased at the smallest available unit (1x1mL). The use of these columns also carries costs associated with acquisition and maintenance of an FPLC machine (capital equipment).

**Figure 2.**
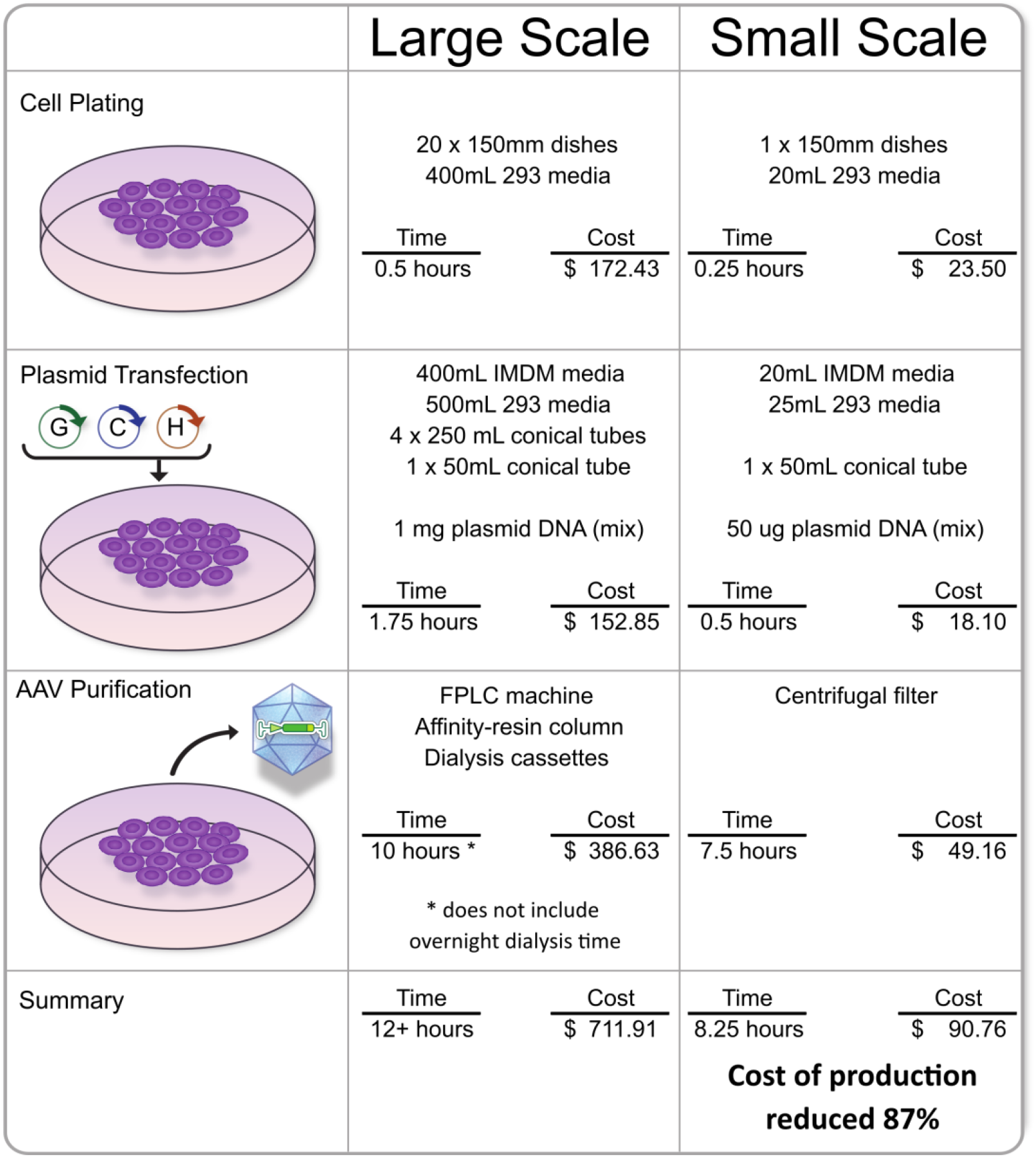
Analysis of reagents and costs for AAV production. A brief comparison of resources needed to package AAV using our established “large scale” protocol and our novel “small scale” protocol. This list does not include cost of plasmid preps, since the pHelper and RepCap plasmids can be more cost-effective when produced at scales larger than “one prep’s worth”. Other labware (i.e., tips, tubes, etc.) are not listed on this table, but were factored into the cost. Total time does not account for incubation of cells between phases or the additional time added by dialysis.

We sought to remove the costly affinity purification step but still provide some mechanism for buffer exchange and volume reduction after cell lysis. Since AAV particles are nearly spherical with a diameter of 26nm and are composed of 60 viral proteins totaling nearly 4 million Daltons (Xie et al., 2002), they are expected to be retained on 100 kilodalton molecular weight cut off (MWCO) spin columns. These filters are available in various load volumes, and the 15mL model was ideally suited to handle the lysate produced from a single 150mm plate.

To test this, we prepared the same AAV genome vector (pAAV EF1a EGFP-KASH, Fig 3A) as either AAV1 or AAV9, using our large scale method of transfecting 20x150mm dishes followed by FPLC/affinity purification and using our prototype small scale method which transfects a single 150mm dish followed by membrane-based filtration, buffer exchange, and concentration (volume reduction). Much of the cell plating and transfection remained similar in both protocols, except that the large scale method prepares the DNA/buffer mixes in four batches each sufficient to transfect stack of five plates, whereas the small scale method prepares these components in a single batch for a single plate. Similar scaled-down volume-based changes were made as needed throughout the transfection and cell lysis steps.

**Figure 3.**
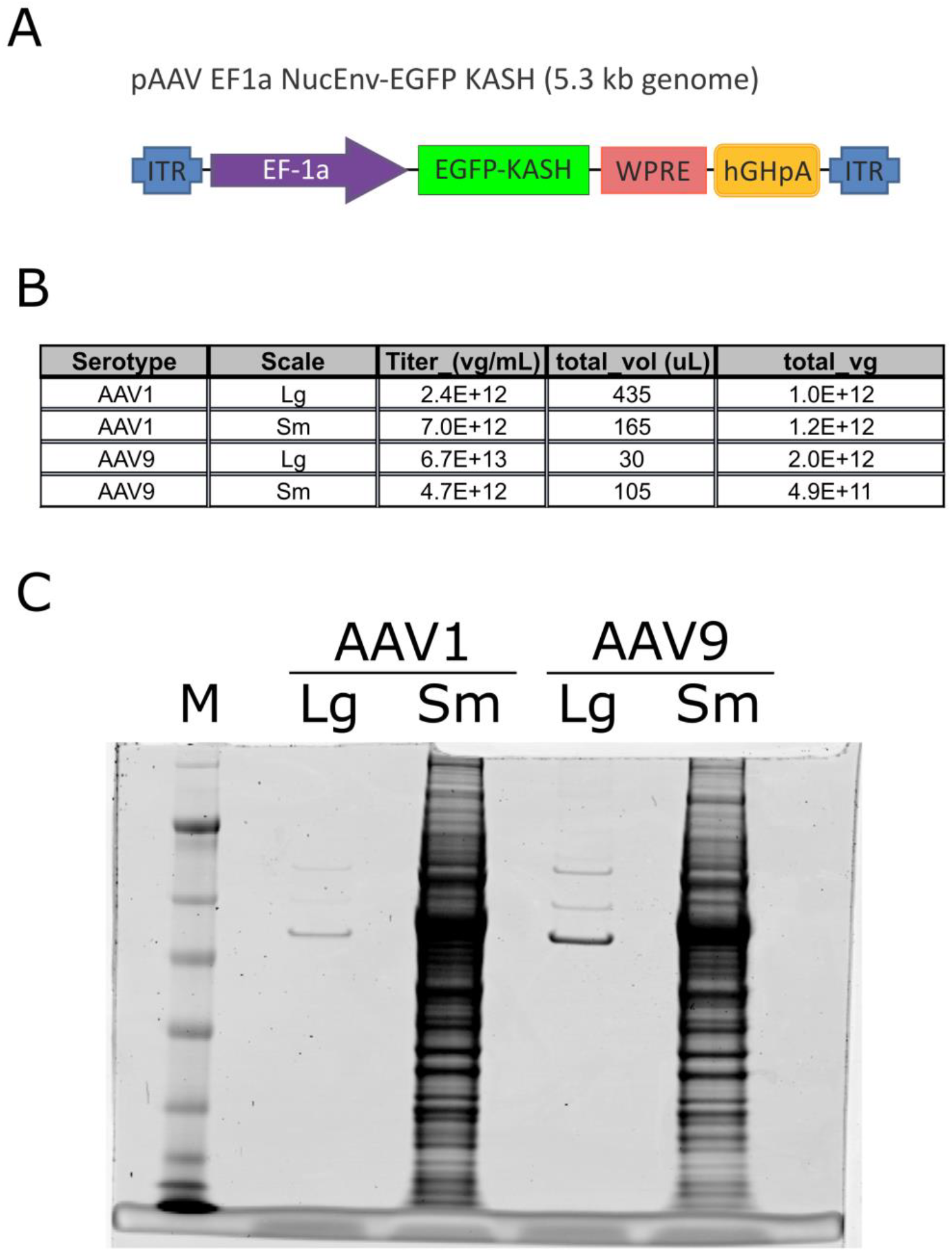
Vector-matched packaging with both protocols. A) A schematic of the AAV vector genome which expresses a GFP-KASH reporter gene under the EF1a promoter. B) The titer and total yield (viral genomes) from the four packaging attempts, as quantified by digital PCR. C) A SimplyBlue-stained NuPAGE gel loaded with the four different virus preps (normalized by titer). The protein size standard is marked “M”.

In summary, a single 150 mm dish produced 5mL of transfected cell lysate that underwent volume reduction to 150uL. The retentate was diluted in 5mL of AAV dialysis buffer and reconcentrated three times. The final retentate of 150uL was then aliquoted for quality control assays and transduction testing.

### Section 2. Comparison of titer and purity

We analyzed the end products from these packaging protocols for viral titer using digital PCR (Fig 3B) and for protein composition by polyacrylamide gel electrophoresis (PAGE) and SimplyBlue staining (Fig 3C). We also tested each lot for bacteria or fungal contamination, and all lots were found to be “negative” (not shown). The large scale preps showed the expected patterns of VP1, VP2, and VP3 on the SimplyBlue-stained gel, whereas the small scale preps were more “complex”, which was expected as they are essentially “whole cell lysates”.

### Section 3 . Comparison of transduction in vitro

The four lots were normalized by titer to 2.4E12 vg/mL. We transduced rat primary cortical neurons (PCNs) with different doses and analyzed viability and expression of the fluorescent reporter nine days post transduction (Fig 4A). The expression levels were varied significantly across the different samples, but all samples showed GFP expression indicating that functionally transducing particles were produced by both workflows (Fig 4B). Large scale AAV1 (left column) shows some GFP puncta, which we associate with overexpression of the EGFP-KASH reporter. These puncta were not present in the viral aliquots (not shown). These puncta were not observed in the large scale AAV9 samples, which showed less GFP expression than the dose matched AAV1. We attribute this to AAV9 have a reduced efficiency for transducing PCNs compared to AAV1.

**Figure 4.**
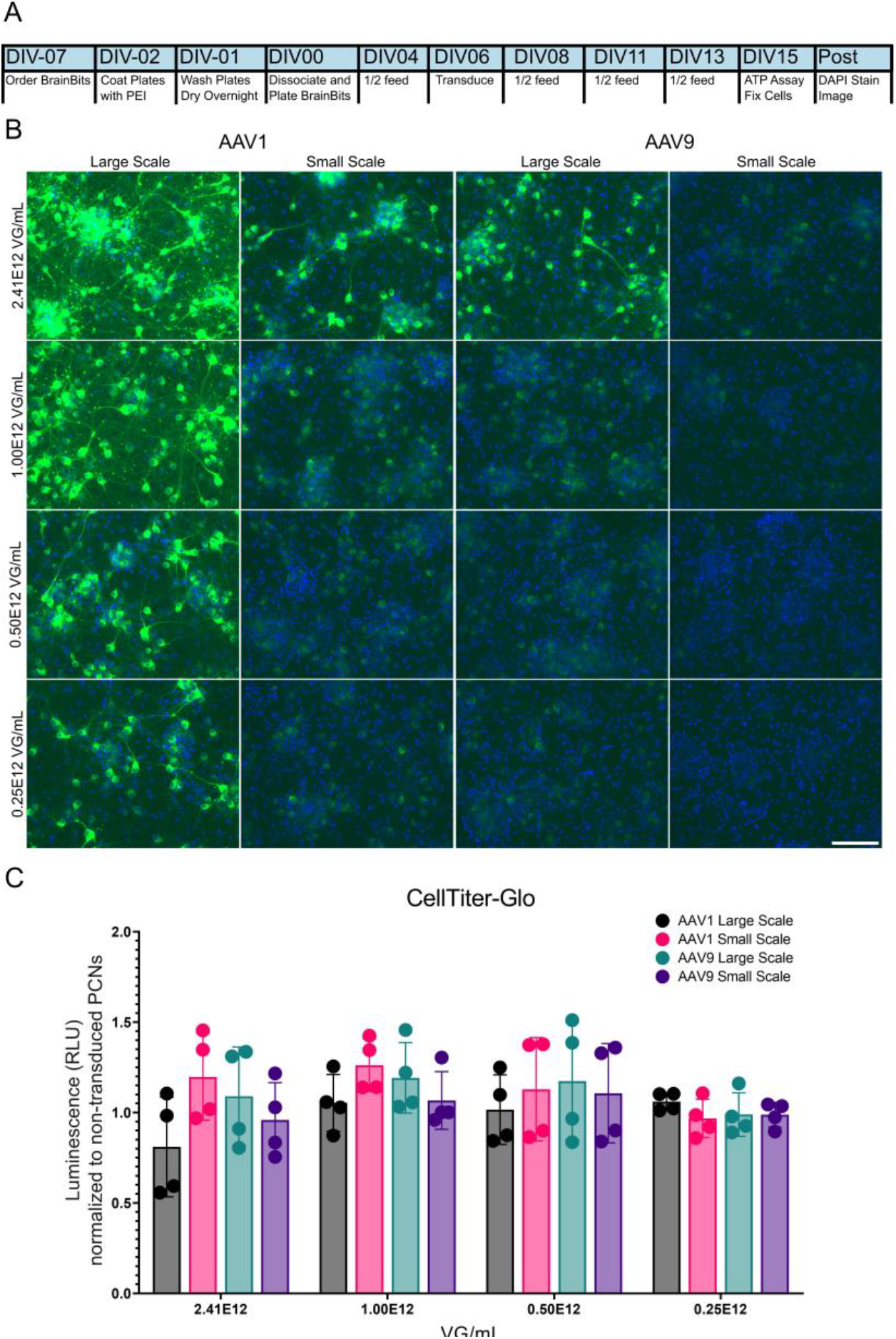
Comparison of vectors using In vitro transduction. (A) A timeline describing the isolation of primary cortical neurons from rat brain tissue, transduction with AAV on DIV6, and paraformaldehyde fixation on DIV15. DIV = days in vitro. (B) A panel of epifluorescent images showing the expression of EGFP-KASH and genomic DNA as stained by DAPI in cells transduced with serial dilutions of AAV. Scale bar is equivalent to 125 microns. (C) A histogram showing the results of a CellTiter-Glo assay to determine cell viability run on DIV15. No significant difference was found using an ordinary two-way ANOVA.

Samples transduced with small scale preps showed less GFP expression than the dose-matched large scale of the same serotype, indicating that the small scale preps may have a reduced transduction efficiency despite have similar titers by digital PCR. Cell health/metabolism as assayed by Promega’s CellTiter-Glo kit showed no differences between transduced and non-transduced cells, for any serotype or viral dose (Fig 4C).

### Section 4. Comparison of transduction in vivo

The four lots were normalized by titer and injected into the striatum of adult male rats. After a recovery period of 14 days, the animals were perfused with PFA and sectioned for immunofluorescence analysis.

At lower magnification, the GFP expression can be observed centered around the injection site for both serotypes and prep scales (Fig 5). Similar to the results obtained from the in vitro transduction, the sides injected with the small scale preps showed less GFP expression than the side injected with the dose-matched large scale of the same serotype. However, we observed that the side injected with large scale AAV9 showed more GFP expression compared to the large scale AAV1. This may be due to the differences in tropism for these two serotypes in the context of primary cortical neurons in culture vs the adult rat brain. At this magnification, we did not observe any differences between serotype or prep-type with regards to tissue necrosis, abscess, infection, etc. We also failed to detect any differences in the staining patterns of GFAP and Iba1, which serve as a marker for gliosis and microglial activation under high magnification (Fig 6)

**Figure 5.**
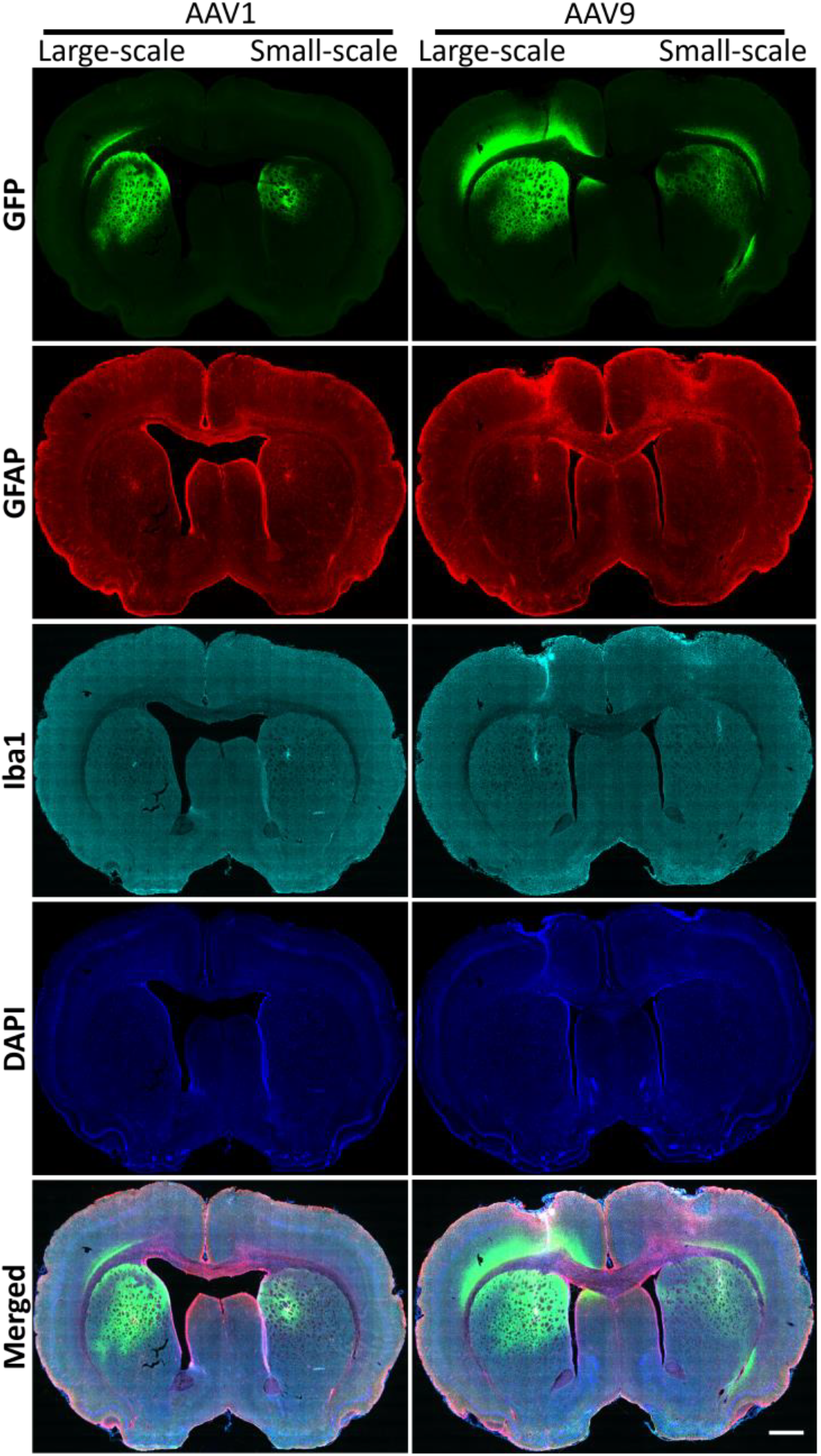
Small scale packaged AAVs do not show increased Iba1 or GFAP staining in striatal tissue. Rats were bilaterally injected in striatum with AAV1 or AAV9 packaged using either large scale or small scale protocols. Tissue sections were stained for GFAP and Iba1. Images of whole sections were captured at 20X magnification. Scale bar 1 mm.

**Figure 6.**
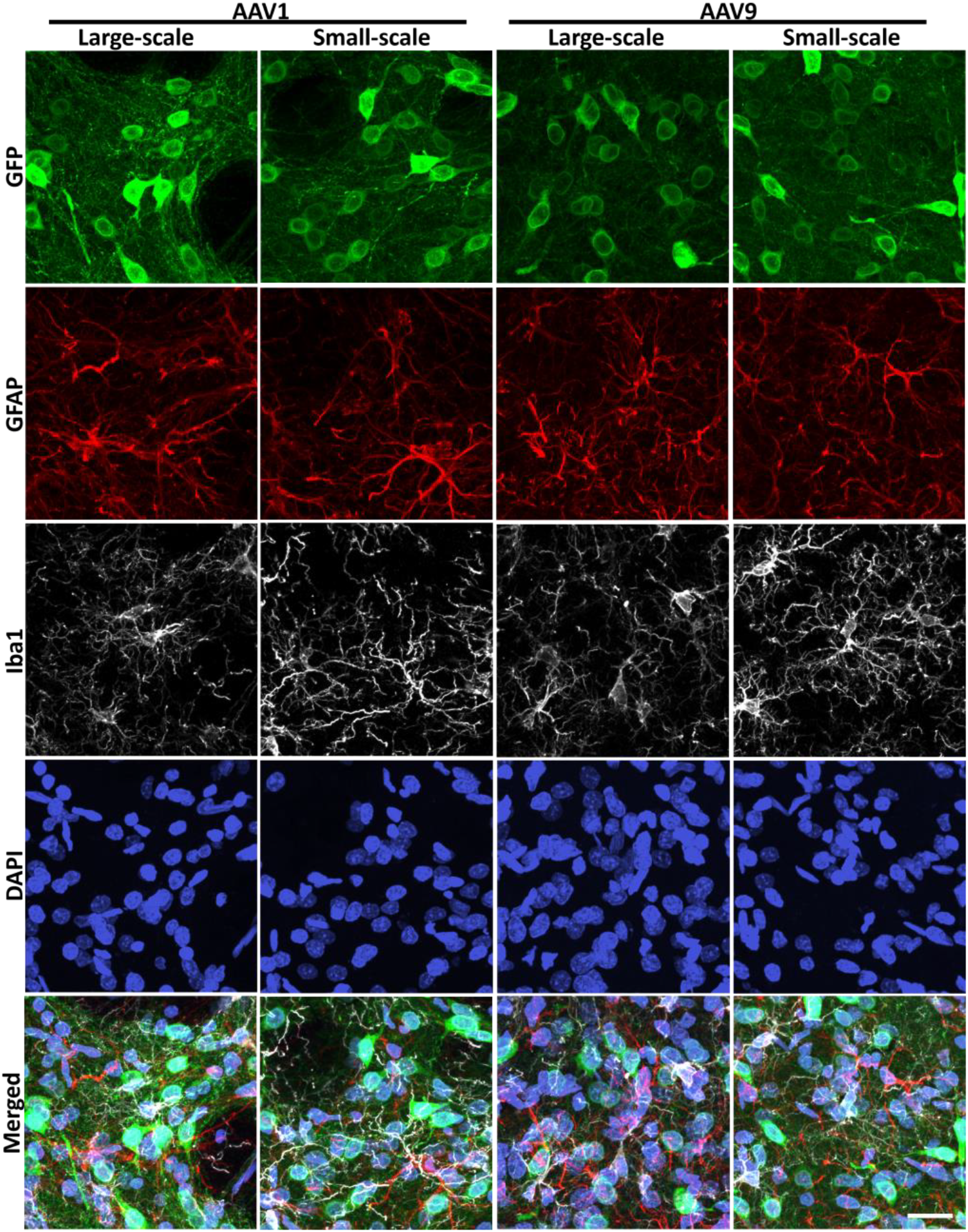
Staining pattern of Iba1 and GFAP is not altered between small-scale packaged and conventionally packaged AAVs. Rats were bilaterally injected in striatum with AAV1 or AAV9 packaged using either large scale or small scale protocols. Tissue sections were stained for GFAP and Iba1. Images were captured at 40X magnification. Scale bar 20 µm.

## DISCUSSION

We set out to develop a workflow for preparing adeno-associated viral vectors that would lower the cost per prep even if that meant accepting a lower yield and purity compared to our standard large scale workflow. Our secondary objective was to increase the throughput of production. In principle, these kinds of improvement would allow us to move forward with more experimental viral genome constructs per project without burdening our research budget or production queue.

By replacing the affinity chromatography step with a centrifugal filter and scaling down the number of 150 mm plates, we were able to produce functional viral particles of AAV1 and AAV9 of sufficient titer for testing in vitro (rat primary cortical neurons) and in vivo (adult rats). These modifications to the production workflow removed the single most expensive consumable (the affinity column) and the longest bottleneck (running the FPLC, one vector at a time). While it may be possible to reuse chromatography columns by stripping them with harsh solvents, we avoid this practice when switching viral genomes due to previous incidents where initial runs of AAV-Cre led to contamination of subsequent preps with unexpected Cre recombination activity.

Moving to a single 15 mL centrifugal filtration device and a single plate transfection had several benefits. First, we can process six tubes corresponding to 6 different viruses in parallel which allows for a substantial increase in the maximum number of different vectors that can be prepped at a time. Preparing six differently transfected plates in parallel does not add a significant burden in terms of incubator space, and the relative amount of viral genome plasmid needed for the transfection can be accommodated by smaller plasmid preparations (midiprep instead of maxiprep).

The AAV vectors produced by the small scale method were able to transduce rat PCNs and led to expression of the GFP reporter. This use case is significant when prototyping viral genome constructs that make use of tissue-specific promoters that restrict transgene expression to neurons (or other cell types), or express transgenes that require specific cellular processes (i.e., synaptic vesicle packaging, neuronal firing, etc.). These kinds of parameters are not always testable by standard transfection of the viral genome plasmid into a transformed cell culture model (HEK293, HeLa, etc.).

Our method allows for the expansive exploration of AAV variant design space, enabling a “fail fast” mindset where prototypes can be tested for efficacy quickly and cheaply in the relevant model before incurring the penalties associated with large scale AAV production (DiPiro and Chisholm-Burns, 2013; McGrath, 2011). For example, the method may be particularly advantageous when testing guide RNA delivery (AAV-CRISPR) or transgene regulation systems (promoters, aptamers, recombination schemes, etc.).

## Supporting information

Sup Doc 1 Large scale AAV packaging

Sup Doc 2 Small scale AAV packaging

## ACKNOWLEDGMENTS

This research was supported by the Intramural Research Program of the NIH, NIDA. AAV viral vectors were produced by the National Institute on Drug Abuse Genetic Engineering and Viral Vector Core Facility (RRID:SCR_022969). We would like to thank Lauren Brick at NIDA Visual Media for assistance with figure generation and graphic design.

---

**Supplemental Table 1.**
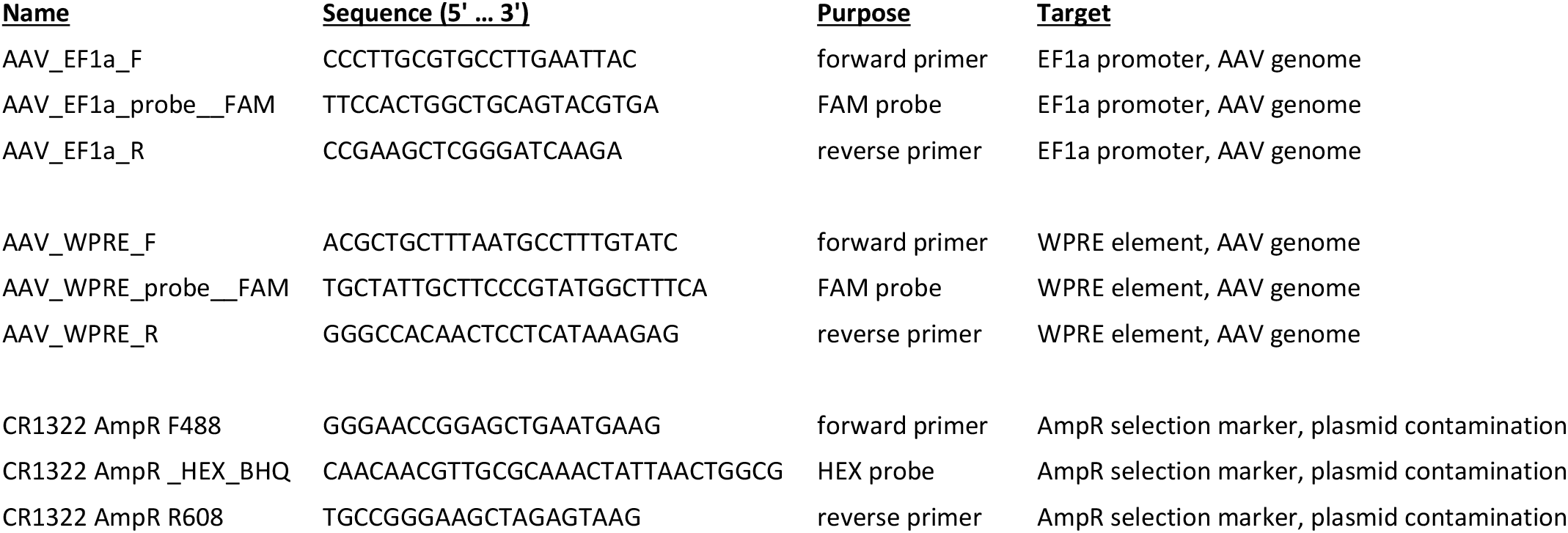
Primers and probes used for titering AAV by digital PCR.

## Notes

### Competing Interest Statement

The authors have declared no competing interest.

