## Supplementary material for "A Rapid Method for Producing Adeno-Associated Viral Vectors Suitable for Transducing Rodent Neurons *in vitro* and *in vivo*": Sup Doc 1 Large scale AAV packaging

### Purpose:

To be able to package, purify, and titer AAV produced from 20 X 150mm dishes of 293XX cells.

### Background

### Procedure

#### Cell Plating

1. Harvest the 293XX cells by aspirating media then rinsing with 10mL 1X PBS .
2. Aspirate 1X PBS and then add 5mL 0.05% Trypsin-EDTA.
3. Place plate back in 37°C incubator for 2-5 minutes. Cells should be visibly lifted from bottom of the plate before moving to the next step.
4. Knock the cells off of plate with 10mL **293XX media** and transfer cell suspension to a 50mL conical tube.
5. Centrifuge cells at 1000RPM for 5 minutes at room temperature.
6. Aspirate supernatant (SN) and resuspend cells in 45ml of **293XX media**.
7. Count live and dead cells. We use an Invitrogen Countess assay and average the results of the two readings. Record all information on the AAV packaging log sheet.
8. Dilute/adjust the cell concentration (based on passage number of cells, see footnotes) to achieve a 70%-85% confluency on the day of transfection (DIV 4).
9. Set up 20 x 150mm plates with 19 ml of **293XX media**. Transfer 1mL of the adjusted cell suspension into each plate. Distribute the cells evenly by gently sliding the plate back and forth and side to side. Do not "swirl" plates.
10. Place cells in incubator. Let cells grow for 4 days (DIV 4) without a media change.
11. OPTIONAL: On DIV 3, thaw 45mL of **2X HBS** per 20 plates of 293XX cells in the refrigerator.

#### Cell Transfection

12. Check cell density by inspecting plates with brightfield microscopy. 75-85% confluent is ideal. If the plates are 50% confluent or less, the transfection should be delayed 24 hours and then assessed again.
13. Image the first plate of each stack of 5 plates under a bright field microscope.
14. Rinse a 25mL burette with dH2O.
15. Fill the burette with **70% isopropanol** and let sit outside the hood until step 18.
16. Working with one stack of five plates at a time, aspirate the media from each plate and replace with 20 mL **Transfection media**, then place back in incubator.
17. Clean the hood
18. Empty and spray the burette, and place in the hood.
19. Sterilize the hood with UV lights for 10-15 minutes.
20. Mix the **2xHBS** by shaking or vortexing.
21. Close the stopcock on the burette. Load burette with 5ml of **2xHBS**. Open stopcock and allow its contents to be passed into a 50mL conical tube.

#### Preparation of Transfection mix (per stack of 5 plates)

22. Prepare “transfection mix” by dispensing 10ml of **0.25M $CaCl_2$** into a new 50ml conical tube.
23. Add the following plasmids to the 50ml conical tube:

    125ug helper plasmid (pHelper, Stratagene)
    35ug capsid plasmid (pAAV2-retro, pXR5, pAAV1, pAAV6, etc.)
    90ug packaging plasmid (contains ITRs and the genetic payload)

24. Vortex the tube briefly on a setting between 4 and 5. Adjust vortex setting as needed to prevent the solution from escaping the tube.
25. Close stopcock on the burette and load with 10ml of **2xHBS**.
26. Dropwise add 10 ml of **2X HBS** to the 10 ml of CaCl2/DNA mixture while constantly vortexing (at a setting between 4-5) Adjust vortex setting as needed to prevent the solution from escaping the tube. The rate of addition is critical; keep the drip slow and make sure there aren’t any signs of precipitation. If precipitation forms, slow the rate of drip.
27. Let this “transfection mix” solution sit at room temp for 5 minutes. (It is possible to set up another solution for 5 plates while waiting for the 5 minutes).

#### Application of transfection mix

28. Remove one stack of 5 plates from the incubator.
29. Tritrate the transfection mix 4 times.
30. Add 4 ml of transfection mix to each plate in a stack of five, while gently rocking media in plate. Add solution gently.
31. Repeat step 29 and 30 for remaining plates of the stack. When finished return the stack of five plates to the 37$^{o}$C incubator.
32. Repeat steps 22-31as needed for the remaining stacks of five plates.

#### Media change

33. 14-16 hours post transfection, aspirate the media from each stack of five plates, and replace with **293XX media** (25mL/plate).

#### AAV Collection

34. 40 hours post-transfection, image top plate of each stack. Acquire brightfield (BF) image, as well as any epifluorescent images that would be relevant/useful to monitor transfection efficiency (i.e. a GFP reporter).
35. Detach the cell monolayer by gently scraping the bottom of each plate with cell scraper.
36. Transfer the cells and media into a 250ml conical tube using a 25mL pipette (5 plates x 25 mL / plate = 125 mL cell suspension / tube).
37. Repeat scraping and collection for each stack of plates.
38. Centrifuge cell suspension for 5 minutes at 2000 rpm, 4˚C.
39. Aspirate and discard the supernatant (SN) with aspirator.
40. Resuspend each pellet in 5mL **AAV suspension buffer** (1mL/plate).
41. Pool the resuspended pellets into a new labeled 50ml conical tube. Volume should be around 25ml at this point. Store in -80$^{o}$C freezer.

#### AAV Purification

##### Lyse cells by freeze/thaw method

42. Thaw cell pellets in 37°C water bath.
43. Freeze cells in a dry ice/ethanol slurry for at least 20 minutes (25 minutes for cells and media).
44. Repeat steps 38 and 39 until cell pellets have undergone a total of 3 freezes (including the initial freezing in the -80°C freezer).
45. Add 50uL 1M MgCl2 (2mM final) and SAN HQ nuclease (to get 400 U/mL final) to each tube of cell lysate and digest at 37°C for 1 hour with shaking.
    **Pro tip:** Set up a 37C incubator with a rotator equipped with a 2 X 50mL flipper rack and tape tube into rack. Turn rack on its side and place on shaker in incubator.
46. Centrifuge digested cell lysate at 3750rpm using Beckman Allegra 6R centrifuge with a 6H-3.8 rotor (or equivalent) for 20 minutes at 4°C.
47. Set up a sterile 100mL glass media bottle with a 0.45um pore size 500mL bottle top filter.
48. Transfer 50mL **1X PBS with 2mM MgCl2** into filter, without applying vacuum, and set it aside.
49. Transfer supernatant (SN) of cell lysates to a new 50 mL conical tube. Discard pellet in 10% bleach.
50. Add 25mL **1X PBS with 2mM MgCl2** to supernatant tube.
51. Vortex the now diluted supernatant tube.
52. Filter diliuted supernatant through a 5.0µm syringe filter using a 50mL syringe directly into the prepared 0.45um bottle top filter containing the **1X PBS with 2mM MgCl2** (from step 48) and apply vacuum.
53. Filter the 0.45um filtrate from previous step through a 0.22µm bottle top filter into a new clean 100mL glass bottle.
54. Reserve the 100mL of 0.22um filtered lysate for use as the "LOAD" in the subsequent FPLC purification section.

##### FPLC Purification using AVB Column (works for AAV1, AAV2, AAV2-retro, AAV6, and AAV8)

Our experience is based heavily on purification of serotype AAV1 with Cytiva AVB columns using an AKTA FPLC machine. For other serotypes and columns (i.e., AAV9), see Footnotes.

We anticipate that users will already have experience with programming, using, and maintaining an FPLC machine. We also assume that the user will begin with an FPLC machine that has been cleaned, rinsed, and primed before moving forward with this section.

##### Purifying the viral particles

55. Install the column (1ml column bed, AVB, Cytiva) into the FPLC machine.
56. Wash column with sterile reverse osmosis (RO) water to remove storage solution (20 column volumes, 2ml/min).
57. Equilibrate column **1xPBS + 2mM MgCl2** (20 column volumes, 2ml/min).
58. Load column with filtered lysate (entire lysate, 2mL/min).
59. Wash column with **1X PBS + 2mM** $\mathbf{MgCl_2}$ (30 column volumes, 2ml/min).
60. Ready the fraction collection system.
61. Elute column with **15mM sodium citrate, pH2.0.** (7.5 column volumes, 1ml/min). Collect 0.7 mL fractions into sterile 1.5 mL tubes.
62. Choose which fractions to move forward by analyzing the UV absorbance peak on the chromatogram to identify which fractions contain the eluted viral particles. Move these fractions forward in the Dialysis section that follows below.

##### Stripping and storing the column

63. Strip column with **15 mM sodium citrate, pH 2.0 + 1M NaCl** (15 column volumes, 2ml/min).
64. Wash column with **1X PBS + 2mM $MgCl_2$** (15 column volume, 2ml/min).
65. Wash column with water (15 column volumes, 2ml/min).
66. Remove column from FPLC machine and set aside.
67. Clean FPLC machine as per manufacturer's instructions.
68. Reinstall column into FPLC machine.
69. Wash column with 20% ethanol. (10 column volumes, 2ml/min).
70. Remove column, store at 4C, for future use.
71. Shutdown FPLC machine as per manufacturer's instructions.

##### Dialysis:

72. Pre-rinse dialysis cassette (1 cassette per fraction) in 1L of **AAV Dialysis buffer** in a 2L beaker.
73. Using a 1mL syringe with an 18g needle, transfer each fraction to separate pre-rinsed dialysis cassette.
74. Label cassettes with fraction number and place in **AAV Dialysis buffer** with a stir bar. Place container in $4^oC$ cabinet on a stir plate with a setting of 60rpm for at least 2 hours (4 hours or more is preferable).
75. After 4 hours of dialysis, replace spent/used **AAV Dialysis buffer** with fresh **AAV Dialysis buffer** buffer and let go overnight. Change dialysis buffer in the morning.
76. After 3rd dialysis for at least 2 hours, remove dialysis cassettes from the **AAV Dialysis buffer** and place in the tissue culture hood.
77. Using a 1mL syringe with an 18ga needle, inject 0.5mL of air into the dialysis cassettes (and leave 0.2mL air in syringe).
78. Turn dialysis cassette upside (with needle still in) and remove the AAV-containing solution from inside the dialysis cassette. Transfer syringe contents to a labeled 1.5mL tube.
79. Transfer 12µL of dialysed virus to a low binding 0.5mL tube.
80. Repeat steps 77-79 for all fractions that were dialyzed.
81. Place 1.5mL tubes containing the dialysed AAV solution in the refrigerator until you are ready to aliquot them for long-term storage. Use the 0.5mL tube(s) containing 12uL of dialyzed AAV (step 74) as samples to analyze vector purity using the "SimplyBlue Safe Stain microwave protocol" in the Footnotes, or equivalent total protein visualization assay.
82. Based on the protein gel, determine which of the collected fraction(s) have the best purity and yield. We generally favor higher concentration over higher volume, and therefore we chose to move forward with a single fraction with the most intense bands on the SimplyBlue gel. Depending on your use-case, you may want to combine one or more fractions into a single tube to get more volume.
83. Dispense the chosen fraction(s) into 0.5mL LoBind tubes (15uL/tube). Store at $-80^oC$.

#### AAV Titering by digital PCR

##### Preparing working dilutions of AAV

84. Thaw a viral aliquot on wet ice. Vortex for 30 sec and spin briefly.
85. Transfer 5uL of thawed virus to a new lobind tube labeled "$10^{-2}$" containing 495uL 1xPBS.
86. Transfer 5uL of $10^{-2}$ virus to a new lobind tube labeled "$10^{-4}$" containing 495uL 1xPBS.
87. Transfer 5uL of $10^{-4}$ virus to a new lobind tube labeled "$10^{-6}$" containing 495uL 1xPBS.

88. Transfer 50uL of $10^{-6}$ virus to a new lobind tube labeled "$10^{-7}$" containing 450uL 1xPBS.
89. Keep all "dilution" tubes at 4C or on ice until ready to add to set up the PCR reactions.

#### dPCR setup

90. Set up dPCR reactions for the $10^{-6}$ and $10^{-7}$ dilutions in triplicate. Use 5uL of diluted AAV in a 15uL dPCR reaction. We use probe-based assays for a sequence expected to be in the viral genome (i.e., the EF1a promoter, WPRE, EGFP, etc.) to quantify viral genomes and an assay expected to be in the plasmid backbone (AmpR) to monitor off-target packaging events and/or plasmid contamination. For further details on dPCR, see footnotes.
91. When dPCR run is complete, get the averages of the triplicate reactions. Then calculate the titer by using this equation:

    Undiluted AAV concentration (vg/mL) = average (dPCR result, templates / uL) * (reaction volume/sample volume) * 1/(sample dilution) * (1000uL/1mL)

    You can then average the calculated undiluted titers for the $10^{-6}$ and $10^{-7}$ reactions, unless one of them is invalid/out of range.

#### Equipment

This is a list of equipment used in this protocol.

| Table. Equipment used in this protocol | |
|---|---|
| **General Description** | **Sourcing information (when available)** |
| Humidified Incubator for Tissue Culture | |
| Fast Protein Liquid Chromatography (FPLC) machine | AKTA Pure FPLC machine |
| Clinical Swinging Bucket Centrifuge | |
| 4C Fridge | |
| Tissue Culture Hood | |
| Invitrogen Countess | |
| Beckman Allegra 6R centrifuge with a 6H-3.8 rotor | |
| Vortexer | |
| 25 mL burette | |
| -20C Freezer | |
| -80C Freezer | |
| Digital PCR (dPCR) machine | |

#### Solutions

This is a list of reagents used in this protocol.

**293XX media:** DMEM High Glucose + 5% Bovine Growth Serum (BGS) + 1% Penicillin-Streptomycin. Sterilized by filtering through a 0.22um bottle top filter.

**Transfection media:** IMDM + 5% BGS Filter through 0.22um bottle top filter to sterilize.

**2X HEPES Buffered Saline (2xHBS):** 56mL 5M NaCl (0.28M final) + 11.9g HEPES (*N*-2-hydroxyethylpiperazine-*N′*-2-ethanesulfonic acid; 0.05 M final) + 3mL 0.5M $Na_2HPO_4$ (1.5mM final) + $ddH_2O$ to 900mL. Titrate to pH 7.0 then add $ddH_2O$ to 1L. Filter through 0.22um bottle top filter to sterilize. Repeat 3 times with pH 7.05, 7.09, and 7.12. Follow packaging transfection steps with a plasmid that has a convenient reporter (5 plates/pH). Determine which specific pH(s) produce the best transfection efficiency based on the reporter expression, and then aliquot those pH(s) into 45mL aliquots and freeze in -20$^o$C.

**0.25M $CaCl_2$:** 36.75g + $ddH_2O$ to 1L. Filter through 0.22um bottle top filter to sterilize.

**AAV suspension buffer:** 50mL 1M Tris-HCL pH 8.0 (50mM final) + 30mL 5M NaCl (150mM final) + 2mL 1M $MgCl_2$ (2mM final) + $ddH_2O$ to 1L. Filter through 0.22um bottle top filter to sterilize.

**1X PBS + 2mM $MgCl_2$:** 100mL 10X PBS + 2mL 1M $MgCl_2$ + $ddH_2O$ to 1L. Filter through 0.22um bottle top filter to sterilize. (Sigma, M1028-10X1ML)

**15mM sodium citrate, pH2.0:** 2.2g Sodium Citrate + ddH2O to 500mL. pH to 2.0 using concentrated HCl. Filter through a nylon 0.22um bottle top filter.

**15mM sodium citrate, pH2.0 + 1M NaCl:** 2.2g Sodium Citrate + 100mL 5M NaCl + ddH2O to 500mL. pH to 2.0 using HCl. Filter through a nylon 0.22um bottle top filter.

**AAV Dialysis buffer (aka AAV dilution buffer):** 100mL 10X PBS + 0.5mL 1M $MgCl_2$ + $ddH_2O$ to 1L

**200mM Citrate + 1M NaCl:** 29.24g Sodium Citrate + 100mL 5M NaCl + ddH2O to 500mL. pH to 2.0 using HCl. Filter through a nylon 0.22um bottle top filter.

**200mM Citrate + 0.4M NaCl:** 29.24g Sodium Citrate + 40mL 5M NaCl + ddH2O to 500mL. pH to 2.0 using HCl. Filter through a nylon 0.22um bottle top filter.

#### Reagents sourcing information:

| Description | Vendor | Catalog Number |
|---|---|---|
| 1X PBS | Life Technologies | 1001004 |
| 0.05% Trypsin-EDTA | Life Technologies | 25300054 |
| AVB column | Global Life Sciences | 28411211 |
| 250ml conical tube | Fisher | 05-538-53 |
| low binding 0.5mL tube | Fisher | 07-200-183 |
| cell scraper | Fisher | 07-200-364 |
| 150mm plates | Fisher | 08-772-6 |
| using a 50mL syringe | Fisher | 13-689-8 |
| 1mL syringe | Fisher | 14-823-30 |
| 18g needle | Fisher | 14-826-5D |
| 50mL conical tube | Starstedt | 62.547.100 |
| 2-Mercaptoethanol | Sigma | 63689-25ML-F |
| SAN HQ (nuclease) | Arcticzymes | 70920-150 |
| 25mL pipette | Sarstedt | 86.1685.001 |
| 70% isopropanol | Fisher | A459-1 |
| CaCl2 | Sigma | C3306-100G |
| 0.45um pore size 500mL bottle top filter | Fisher | FB12566511 |
| See Blue Plus 2 | Life Technologies | LC5925 |
| Simply Blue SafeStain | Life Technologies | LC6060 |
| MOPS buffer | Life Technologies | NP0001 |
| 4X loading dye | Life Technologies | NP0007 |
| 10-well Bis-Tris gel | Life Technologies | NP0321BOX |
| Slide-A-Lyzer dialysis cassette (0.1-.05mL 10,000 MWCO) | Fisher | PI66383 |
| 5.0µm syringe filter | EMD Millipore | SLV025LS |
| NucleoBond® Xtra Maxi Plus | Takara Bio USA | 740416.1 |

#### Footnotes

##### Plasmids
We prepare our plasmids using Macherey-Nagel NucleoBond® Xtra Maxi Plus.  All plasmids should be dissolved in endotoxin-free water or 1xTE and should have concentrations above 500ng/uL.

##### 293XX cells
293XX cells are a derivative of HEK293 that NIDA GEVVC obtained Xiao Xiao (UPITT), via Brandon Harvey (NIDA).  We continue to use this cell line for consistency across production runs, but this protocol should work with any adherent HEK293-related cell line.

For routine maintenance of this cell line, split 1-to-5 every 2-3 days. When setting up for an AAV packaging event, they should be split 1-to-4 into at least two 150mm plates 48 hours before Cell Plating Step 1.

We have observed that 293XX cells grow/divide more quickly as the passage number increases. Therefore, the number of seeded cells needed to reach a proper confluency is passage-number dependent.

| Table 1. 293XX cell passage number to plating density | |
|---|---|
| 293XX Passage number | Cell density for plating (cells / mL) |
| P1-12 | 2E6 |
| P13-P15 | 1.9E6 |
| P16-P18 | 1.8E6 |
| P18-P20 | 1.7E6 |
| P21-P30 | 1.6E6 |
| P31+ | Do not use, thaw an aliquot from lower passage. |

#### Using POROS AA9 Column (AAV9)

The use of POROS AAV9 columns from Thermo will require minor modifications to the solutions used during the FPLC purification process. The steps below will replace the analogous steps previous described for AVB columns (steps 55-71).

##### Purifying the viral particles

55. Install the column (1ml column bed, POROS AAV9, Thermo) into the FPLC machine.
56. Wash column with sterile reverse osmosis (RO) water to remove storage solution (20 column volumes, 2ml/min).
57. Equilibrate column **1xPBS + 2mM MgCl2** (20 column volumes, 2ml/min).
58. Load column with filtered lysate (entire lysate, 2mL/min).
59. Wash column with **1X PBS + 2mM $MgCl_2$** (30 column volumes, 2ml/min).
60. Ready the fraction collection system.
61. Elute column with **200mM sodium citrate, pH2.0 + 0.4M NaCl.** (7.5 column volumes, 1ml/min). Collect 0.7 mL fractions into sterile 1.5 mL tubes.
62. Choose which fractions to move forward by analyzing the UV absorbance peak on the chromatogram to identify which fractions contain the eluted viral particles. Move these fractions forward in the Dialysis section that follows below.

##### Stripping and storing the column

63. Strip column with **200 mM sodium citrate, pH 2.0 + 1M NaCl** (15 column volumes, 2ml/min).
64. Wash column with **1X PBS + 2mM $MgCl_2$** (15 column volume, 2ml/min).
65. Wash column with water (15 column volumes, 2ml/min).
66. Remove column from FPLC machine and set aside.
67. Clean FPLC machine as per manufacturer's instructions.
68. Reinstall column into FPLC machine.
69. Wash column with 20% ethanol. (10 column volumes, 2ml/min).
70. Remove column, store at 4C, for future use.
71. Shutdown FPLC machine as per manufacturer's instructions.

#### SimplyBlue Safe Stain microwave protocol

1. Add 4uL of 4X loading dye with 2-Mercaptoethanol to each 0.5 mL lobind tube containing 12uL of sample.
2. Vortex samples then spin in minifuge for 10 seconds.
3. Heat to 70C for 10 minutes with 650rpm shaking in thermomixer.
4. Prepare 1X MOPS buffer.
5. Run 10uL of See Blue Plus 2 and 16uL of the samples on a 4-20%, 10-well Bis-Tris gel for 50 minutes at 200V.
6. Remove gel from casing into 100mL ultra pure water and microwave for 45 seconds.
7. Place gel on plate rotator for 1 minute at room temp.
8. Remove water and repeat above microwave and shake steps 2 more times (3 total).
9. After gel shakes for last time in water, remove water and add 30mL Simply Blue SafeStain and microwave for 40 seconds.
10. Shake on belly dancer (setting of 4) for 5 minutes at room temp.
11. Discard Simply Blue Safe Stain in liquid waste carboy and rinse with DI water.
12. Add 100mL ultra pure water and place on belly dancer for 10 minutes at room temp.
13. Image gel using Azure imager.
14. Based on the protein gel, determine which of the collected fraction(s) have the best purity and yield. We generally favor higher concentration over higher volume, and therefore we chose to move forward with a single fraction with the most intense bands on the SimplyBlue gel. Depending on your use-case, you may want to combine one or more fractions into a single tube to get more volume.

### AAV Packaging Log Sheet

**Cells Setup**

Date/Time plated: ____________________ Cell Passage #: _____ Initials: ______________

| Countess | Live Cells | Dead Cells | % Viable |
|---|---|---|---|
| **Section A** | | | |
| **Section B** | | | |
| **Average** | | | |

Cell concentration and volume per plate:__________________________________________

**Transfection:**

Date/Time: ____________________ Virus Lot#: _____________________ Initials: ____________

2XHBS pH: __________ 2XHBS Prep date: __________ CaCl2 Prep date: __________

| | Plasmid | pOTTC # | plasmid DNA prep date | Concentration mg/mL | Volume (uL) |
|---|---|---|---|---|---|
| Gene: | _____________ | __________ | ___________ | _____________ | __________ |
| Helper | ______________ | __________ | ___________ | _____________ | ___________ |
| Capsid | ______________ | __________ | ___________ | _____________ | ___________ |

**Media Change**: (Date/Time)______________ **Harvest:** (Date/Time)_______________

**No. of plates:** ________________ **AAV suspension buffer date:**____________

**Notes:**

#### AAV Purification log sheet

**Experiment #:** ______________________________

**Purification Method:** __________________ Thaw 1 start time: __________________

Freeze 2 start time: ____________________ Thaw 2 start time: __________________

Freeze 3 start time: ____________________ Thaw 3 start time: __________________

Volume of cell pellet: ___________________ Endonuclease: ____________________

Company: _____________________________ Lot #: ____________________________

Concentration: _______________________ Volume of Endonuclease: ___________

Stock concentration of $MgCl_2$: ____________ Volume of $MgCl_2$centration: __________

Incubator shake start time: _________________________________________________

**Date/time spun:** ___________________ Temp/RPM/Time spun: ____________

**Buffer Information (Date / Lot #)** PBS + 2mM $MgCl_2$: ________________

15mM Sodium Citrate: ___________________________________________

15mM Sodium Citrate + 1M NaCl: __________________________________

Column # used: ______________________ Time loaded on column: _______________

Fractions Dialyzed: __________________ Date/Time Dialysis 1: _______________

Date/Time Dialysis 2: _________________ Date/Time Dialysis 3: _______________

**Protein gel:**

Date/Time heated: ___________________ Temp/RPM/time heated: _____________

Time ran on gel: ______________ Type of gel: _____________________________

Voltage of run: _______________ Duration of gel run: _______________________

Gel setup:

| Lane | Sample | Label |
|---|---|---|
| 1 | | |
| 2 | | |
| 3 | | |
| 4 | | |

Fraction(s) aliquotted: ____________________________

| AAV Box # | Virus name | Virus lot # | Chromatogram Peak 280/250 | pOTTC # | 15uL aliquots | 50uL aliquots |
|---|---|---|---|---|---|---|

**Notes:**

**Storage site:** ___________________________
