## Supplementary material for "A Rapid Method for Producing Adeno-Associated Viral Vectors Suitable for Transducing Rodent Neurons *in vitro* and *in vivo*": Sup Doc 2 Small scale AAV packaging

### Purpose:

To be able to package, purify, and titer AAV produced from 1 X 150mm dishes of 293XX cells.

### Background

### Procedure

#### Cell Plating

1. Harvest the 293XX cells by aspirating media then rinsing with 10mL 1X PBS .
2. Aspirate 1X PBS and then add 5mL 0.05% Trypsin-EDTA.
3. Place plate back in 37°C incubator for 2-5 minutes. Cells should be visibly lifted from bottom of the plate before moving to the next step.
4. Knock the cells off the plate with 10mL **293XX media** and transfer cell suspension to a 50mL conical tube.
5. Centrifuge cells at 1000RPM for 5 minutes at room temperature.
6. Aspirate supernatant (SN) and resuspend cells in 45ml of **293XX media**.
7. Count live and dead cells. We use an Invitrogen Countess assay and average the results of the two readings. Record all information on the AAV packaging log sheet.
8. Dilute/adjust the cell concentration (based on passage number of cells, see footnotes) to achieve a 70%-85% confluency on the day of transfection (DIV 4).
9. Set up 1 x 150mm plates with 19 ml of **293XX media**. Transfer 1mL of the adjusted cell suspension into each plate. Distribute the cells evenly by gently sliding the plate back and forth and side to side. Do not "swirl" plates.
   Repeat Step 9 for up to six plates (each plate being used for a separate viral packaging)
10. Place cells in incubator. Let cells grow for 4 days (DIV 4) without a media change.
11. OPTIONAL: On DIV 3, thaw 45mL of **2X HBS** per 20 plates of 293XX cells in the refrigerator.

#### Cell Transfection

12. Check cell density by inspecting plates with brightfield microscopy. 75-85% confluent is ideal. If the plates are 50% confluent or less, the transfection should be delayed 24 hours and then assessed again.
13. Image the plate under a bright field microscope.
14. Aspirate the media and replace with 20 mL **Transfection media**, then place back in incubator.
    Repeat Steps 12-14 for all plates being used for packaging.

#### Preparation of Transfection mix (per 1 x 150 mm plate)

15. Mix the **2xHBS** by shaking or vortexing.
16. Prepare "transfection mix" by dispensing 2mL of **0.25M $CaCl_2$** into a new 50ml conical tube.
17. Add the following plasmids to the 50ml conical tube:

| | |
|---|---|
| 25ug | helper plasmid (pHelper, Stratagene) |
| 7ug | capsid plasmid (pAAV2-retro, pXR5, pAAV1, pAAV6, etc.) |

18ug packaging plasmid (contains ITRs and the genetic payload)

18. Vortex the tube briefly on a setting between 4 and 5. Adjust vortex setting as needed to prevent the solution from escaping the tube.
19. Using a 5mL pipet, dropwise add 2ml of 2X HBS to the 2ml of CaCl2/DNA mixture while constantly vortexing (at a setting between 4-5) Adjust vortex setting as needed to prevent the solution from escaping the tube. The rate of addition is critical; keep the drip slow and make sure there aren't any signs of precipitation. If precipitation forms, slow the rate of drip.
20. Let this "transfection mix" solution sit at room temp for 5 minutes. (It is possible to set up another transfection mix for another plate while waiting for the 5 minutes).

##### Application of transfection mix (per 1 x 150 mm plate)

21. Remove the plate from the incubator.
22. Tritrate the transfection mix 4 times.
23. Add the transfection mix to plate, while gently rocking media in plate. Add solution gently.
24. Return the plate to the 37$^{o}$C incubator.
25. Repeat steps 15-24 as needed for the remaining transfection mixes/plates.

##### Media change

26. 14-16 hours post transfection, aspirate the media from each transfected plates, and replace with **293XX media** (25mL/plate).

#### AAV Collection

27. 40 hours post-transfection, image the transfected plate. Acquire brightfield (BF) image, as well as any epifluorescent images that would be relevant/useful to monitor transfection efficiency (i.e., a GFP reporter).
28. Detach the cell monolayer by gently scraping the bottom of each plate with cell scraper.
29. Transfer the cells and media into a 50mL conical tube using a 25mL pipette.
30. Repeat steps 27-29 as needed for the remaining transfected plates.
31. Centrifuge cell suspension for 5 minutes at 2000 rpm, 4˚C using Beckman Allegra 6R centrifuge with a 6H-3.8 rotor (or equivalent).
32. Aspirate and discard the supernatant (SN) with aspirator.
33. Resuspend cell pellet in 5mL **AAV suspension buffer**.
34. Place tube in dry ice/ ethanol slurry.
35. Repeat steps 32-34 as needed for the remaining transfection mixes/plates.

#### AAV Processing

##### Lyse cells by freeze/thaw method

36. Allow cells to stay in dry ice / ethanol slurry for 10 minutes.
37. Thaw cell pellets in 37$^{o}$C water bath for 7 minutes.
38. Repeat steps 36 and 37 until cell pellet(s) have undergone 3 freeze/thaw cycles.
39. Add 10uL 1M MgCl2 (2mM final) and SAN HQ nuclease (to get 400 U/mL final) to each tube of cell lysate and digest at 37$^{o}$C for 1 hour with shaking.
    **Pro tip:** Set up a 37C incubator with a rotator equipped with a 2 X 50mL flipper rack and tape tube into rack. Turn rack on its side and place on shaker in incubator.
40. Centrifuge digested cell lysate at 3750rpm using Beckman Allegra 6R centrifuge with a 6H-3.8 rotor (or equivalent) for 20 minutes at 4$^{o}$C.

41. While spinning, spray off the top loading balance and place in the tissue culture hood.
42. Label a new set of 50mL conical tubes (1 for each AAV being processed).
43. Label one 100,000 MWCO Amicon Ultra-15 centrifugal filter for each AAV being processed.
44. Transfer virus-containing supernatant to a 5mL syringe with a 5.0um syringe filter attached.
45. Filter virus-containing supernatant into the newly labeled 50mL conical tube.
46. Filter the virus-containing supernatant again using the same syringe but now with a 0.45um filter attached.
    Dispense filtrate back into the same 50mL conical tube (step 47).
47. Filter the virus-containing supernatant a third time with using the same syringe but now with a 0.22um syringe filter attached dispensing into one of the labeled centrifugal filters.
48. Balance all tubes under the hood using **AAV dialysis buffer** as balancing solution.
49. Spin all tubes at 4000g, 4$^{o}$C, 40 minutes .
50. Once spin is complete, return tubes to the tissue culture hood and discard flowthrough.
51. Add 5mL of **dialysis buffer** with 5mL pipet.
52. Mix **AAV dialysis buffer** with the retentate by tritrating with a 1mL pipettor at least five times.
53. Repeat steps 48 to 52 .
54. Repeat steps 48 to 49. Once final spin is complete, discard flowthrough and transfer the concentrated AAV to a 0.5mL low binding tube with a P200 pipette.
55. Dispense the concentrated AAV into 0.5mL LoBind tubes (15uL/tube). Store at -80$^{o}$C.

#### AAV Titering by digital PCR

##### Preparing working dilutions of AAV

56. Thaw a viral aliquot on wet ice. Vortex for 30 sec and spin briefly.
57. Transfer 5uL of thawed virus to a new LoBind tube labeled “$10^{-2}$” containing 495uL 1xPBS.
58. Transfer 5uL of $10^{-2}$ virus to a new LoBind tube labeled “$10^{-4}$” containing 495uL 1xPBS.
59. Transfer 50uL of $10^{-4}$ virus to a new LoBind tube labeled “$10^{-5}$” containing 450uL 1xPBS.
60. Transfer 50uL of $10^{-5}$ virus to a new LoBind tube labeled “$10^{-6}$” containing 450uL 1xPBS.
61. Keep all “dilution” tubes at 4C or on ice until ready to add to set up the PCR reactions.

##### dPCR setup

62. Set up dPCR reactions for the $10^{-5}$ and $10^{-6}$ dilutions in triplicate. Use 5uL of diluted AAV in a 15uL dPCR reaction. We use probe-based assays for a sequence expected to be in the viral genome (i.e., the EF1a promoter, WPRE, EGFP, etc.) to quantify viral genomes and an assay expected to be in the plasmid backbone (AmpR) to monitor off-target packaging events and/or plasmid contamination. For further details on dPCR, see footnotes.
63. When dPCR run is complete, get the averages of the triplicate reactions. Then calculate the titer by using this equation:

    Undiluted AAV concentration (vg/mL) = average (dPCR result, templates / uL) * (reaction volume/sample volume) * 1/(sample dilution) * (1000uL/1mL)

    You can then average the calculated undiluted titers for the $10^{-5}$ and $10^{-6}$ reactions, unless one of them is invalid/out of range.

#### Equipment

This is a list of equipment used in this protocol.

Table. Equipment used in this protocol

| **General Description** | **Sourcing information (when available)** |
|---|---|
| Humidified Incubator for Tissue Culture | |
| Fast Protein Liquid Chromatography (FPLC) machine | AKTA Pure FPLC machine |
| Clinical Swinging Bucket Centrifuge | |
| 4C Fridge | |
| Tissue Culture Hood | |
| Invitrogen Countess | |
| Beckman Allegra 6R centrifuge with a 6H-3.8 rotor | |
| Vortexer | |
| 25 mL burette | |
| -20C Freezer | |
| -80C Freezer | |
| Digital PCR (dPCR) machine | |

NIDA GEVVC PROTOCOL – AAV packaging and purification by FPLC, 1 x 150mm plates

#### Solutions

This is a list of reagents used in this protocol.

**293XX media:** DMEM High Glucose + 5% Bovine Growth Serum (BGS) + 1% Penicillin-Streptomycin. Sterilized by filtering through a 0.22um bottle top filter.

**Transfection media:** IMDM + 5% BGS Filter through 0.22um bottle top filter to sterilize.

**2X HEPES Buffered Saline (2xHBS):** 56mL 5M NaCl (0.28M final) + 11.9g HEPES (N-2-hydroxyethylpiperazine-N[illegible]-2-ethanesulfonic acid; 0.05 M final) + 3mL 0.5M Na2HPO4 (1.5mM final) + ddH2O to 900mL. Titrate to pH 7.0 then add ddH2O to 1L. Filter through 0.22um bottle top filter to sterilize. Repeat 3 times with pH 7.05, 7.09, and 7.12. Follow packaging transfection steps with a plasmid that has a convenient reporter (5 plates/pH). Determine which specific pH(s) produce the best transfection efficiency based on the reporter expression, and then aliquot those pH(s) into 45mL aliquots and freeze in -20oC.

**0.25M $CaCl_2$:** 36.75g + $ddH_2O$ to 1L. Filter through 0.22um bottle top filter to sterilize.

**AAV suspension buffer:** 50mL 1M Tris-HCL pH 8.0 (50mM final) + 30mL 5M NaCl (150mM final) + 2mL 1M $MgCl_2$ (2mM final) + $ddH_2O$ to 1L. Filter through 0.22um bottle top filter to sterilize.

**AAV Dialysis buffer (aka AAV dilution buffer):** 100mL 10X PBS + 0.5mL 1M $MgCl_2$ + $ddH_2O$ to 1L

#### Reagents sourcing information:

| Description | Vendor | Catalog Number |
|---|---|---|
| 1X PBS | Life Technologies | 1001004 |
| 0.05% Trypsin-EDTA | Life Technologies | 25300054 |
| 1mL pipette tip | Rainin | 30389212 |
| P200 pipette tip | Rainin | 30389239 |
| low binding 0.5mL tube | Fisher | 07-200-183 |
| cell scraper | Fisher | 07-200-364 |
| 150mm plates | Fisher | 08-772-6 |
| 5mL syringe | Fisher | 14-955-458 |
| 50mL conical tube | Starstedt | 62.547.100 |
| SAN HQ (nuclease) | Arcticzymes | 70920-150 |
| 5mL pipette | Sarstedt | 86.1253.001 |
| 25mL pipette | Sarstedt | 86.1685.001 |
| 70% isopropanol | Fisher | A459-1 |
| CaCl2 | Sigma | C3306-100G |
| 0.45um pore size 500mL bottle top filter | Fisher | FB12566511 |
| 0.22um syringe filter | EMD Millipore | SLGP033RS) |
| 0.45um syringe filter | EMD Millipore | SLHV033RS |

**Notes:**
